# Paracrine Bone-Derived Senescent Secretome Induces Spatially Patterned ECM and Biomechanical Vulnerability in Human Brain Organoids

**DOI:** 10.1101/2025.08.11.669674

**Authors:** Chanul Kim, Luke Daniel Ofria, Anannya Kshirsagar, Elizabeth Gomez Flores, Sai Kulkarni, Alan Y. Liu, James L. Kirkland, Annie Kathuria, Maryam Tilton

**Author notes:** Corresponding authors: M. Tilton and A. Kathuria.

## Abstract

Aging is increasingly recognized as a systemic process, yet the mechanisms by which senescent cells’ signal from peripheral tissues accelerate brain aging remain poorly defined. Here, we used chronic exposure of human cerebral organoids to the secretome of senescent osteocytes to investigate how peripheral aging signals reshape brain tissue architecture. We combined spatially resolved optical fiber-based interferometry nanoindentation with transcriptomic and immunofluorescence profiling, demonstrating that bone-derived senescence-associated secretory phenotype (SASP) factors induce a biphasic mechanical response, early global tissue softening, followed by the emergence of discrete hyper-stiff microdomains. This spatially heterogeneous biomechanical remodeling was accompanied by upregulation of extracellular matrix (ECM), inflammatory, and senescence pathways, and suppression of neurodevelopmental and synaptic gene networks. Our results reveal that chronic paracrine SASP exposure from senescent osteocytes drives localized ECM reorganization and mechanical vulnerability in human brain tissue, providing mechanistic insight into how peripheral cellular senescence may contribute to regional brain fragility during aging.

## Introduction

Aging is a major risk factor for both neurodegenerative diseases, such as Alzheimer’s disease (AD), and musculoskeletal deterioration, including osteoporosis^1–4^. Epidemiology studies indicate that individuals over the age of 65 with bone loss have a 1.2 to 1.4-fold higher risk of developing dementia than those without evident bone loss^1,2^. Since dementia cases are projected to exceed 70 million globally by 2030, and the combined economic burden of bone loss and neurodegeneration is expected to surpass $1.2 trillion by 2050^1,3,5,6^, understanding mechanistic links between bone health and brain aging has become an urgent public health priority^1–4^. While traditionally treated as independent conditions, emerging research suggests that bone and brain aging share common molecular drivers, including chronic inflammation, oxidative stress, and cellular senescence^3,5,6^, highlighting the need to study the “bone-brain axis” holistically. Once considered solely a structural tissue, bone is now recognized as active endocrine and immunoregulatory organ^7,8^. Osteocytes and other bone cells release a broad spectrum of signaling molecules, including hormones, cytokines, and extracellular vesicles, that exert systemic effects and modulate the function of distant organs such as the brain^7,9,10^.

One key mechanism linking bone aging to brain dysfunction is cellular senescence ^3,5,6^. Senescent bone cells, including osteocytes and bone marrow stromal cells (BMSCs), accumulate with age and secrete a pro-inflammatory, tissue-remodeling secretome known as the senescence-associated secretory phenotype (SASP). Canonical SASP factors (e.g., IL-1β, IL-6, TNF-α, matrix metalloproteinases) as well as bone-specific molecules such as sclerostin, which can cross the blood-brain barrier (BBB), are elevated in aged bone^11,12^. These bone-derived factors can disrupt neural homeostasis, promote neuroinflammation, impair neurogenesis, and accelerate cognitive decline^13^. However, despite growing recognition of the systemic influence of bone, the specific impact of bone-derived senescent secretomes on brain aging remains poorly understood.

Existing brain aging models have focused mainly on intrinsic neuronal and glial processes^14^, overlooking systemic contributions from aging peripheral organs. Yet, cross-organ aging signals, reflected in mitochondrial deficits, proteostasis decline, and chronic inflammation, are shared features of aged cells across organs. Furthermore, preclinical studies show that youthful bone-derived factors can enhance memory and systemic rejuvenation^15,16^, underscoring the potential for bone-secreted molecules to modulate brain function.

Here, we developed a novel cross-organ aging model in which patient-derived cerebral organoids were chronically exposed to conditioned media rich in SASP factors (SASP-CM) secreted by senescent bone cells. In this study, human cerebral organoids cultured for 14 days in either background growth media (Control) or the conditioned media collected from irradiation-induced senescent osteocytes (SASP group). The conditioned media were harvested on day 21 post-irradiation to ensure robust accumulation of SASP factors, following validated senescence induction protocols from our team^17–20^. By applying longitudinal micromechanical mapping, live-cell imaging, bulk transcriptomic profiling, and immunofluorescence analyses, we examined how chronic exposure to bone-derived senescent secretomes affects the biomechanical properties, inflammatory activation, and molecular aging signatures of brain organoids. This approach provides a tractable human platform for interrogating systemic propagation of senescence and its impact on brain tissue aging. While this *in vitro* model does not capture the full complexity of the bone-brain axis, it uniquely enables direct, mechanistic investigation of bone-derived senescent signals in a human context. To our knowledge, this is the first study to demonstrate that bone-derived SASP factors can drive neuroinflammatory and aging-associated biomechanical and biomolecular remodeling in human cerebral organoids, establishing a foundation for future work on systemic senescence in brain aging.

## Results & Discussion

### Bone-derived SASP exposure induced subtle but progressive cytoskeletal and viability alterations in cerebral organoids

To determine whether paracrine exposure to bone-derived senescent secretomes (SASP factors) compromises the viability and structural integrity of human cerebral organoids, we performed longitudinal live/dead staining and multiplex immunofluorescence for cytoskeletal and nuclear markers (Fig.1b-c). Live/dead imaging on Day 7 (Fig.1b) revealed that both control and SASP-treated organoids retained high overall viability, as evidenced by robust Calcein-AM (live) staining and only modest EthD-1 (dead) signal. Notably, SASP-treated organoids exhibited a subtle but consistent increase in localized EthD-1-positive clusters, particularly in the interior regions, suggesting microenvironmental vulnerability that may not reach significance at the whole-organoid scale. This observation is consistent with prior report that SASP factors promote selective vulnerability within specific microenvironments of neural tissue, rather than causing uniform cell death throughout the tissue^21^.

Immunofluorescence for F-actin and nuclear morphology across time points (Fig.1c) revealed that, while control organoids maintain well-defined peripheral actin networks and dispersed nuclei at D7, SASP-treated organoids already demonstrate early signs of actin reorganization. By D14, both control and SASP-treated groups exhibit progressive increases in actin filament complexity and nuclear clustering; however, these changes are more pronounced in the SASP-D14 group, which displays fragmented F-actin domains and compacted nuclear regions. This observation suggests that paracrine SASP exposure may accelerate cytoskeletal disarray and tissue compaction, phenotypes increasingly recognized as precursors to loss of neural connectivity and cellular resilience^20,22–24^.

While the overall architecture remains grossly intact at early stages, our results point to a cumulative impact of the bone-derived SASP factors on organoid microarchitecture over time. At D14, the increased heterogeneity in actin organization and nuclear distribution, particularly in SASP-exposed organoids, may signal the emergence of localized vulnerability, potentially priming the tissue for subsequent inflammatory or degenerative cascades. These findings align with recent work demonstrating that chronic SASP exposure can prime neural tissue for stress-induced dysfunction even in the absence of overt cell loss^25^.

While previous studies have implicated systemic SASP factors in promoting neuroinflammation and brain aging^25^, our data provide the first direct evidence that bone-derived senescent secretomes can induce early, spatially restricted disruptions in human brain organoid viability and cytoskeletal integrity. The subtle but progressive nature of these alterations distinguishes SASP-mediated neural remodeling from overt cytotoxicity observed in acute injury models^26,27^, highlighting the unique pathophysiological context of aging-associated cross-organ signaling. The spatial association of cell death foci with cytoskeletal disorganization merits further exploration^28,29^, particularly within microenvironmental stress as a hallmark of early neurodegenerative processes. To directly interrogate the mechanobiological consequences of these localized vulnerabilities, we implemented a live, non-destructive optical fiber-based interferometry nanoindentation platform (Fig.2), enabling high-resolution mapping of regional tissue mechanics in response to bone-derived SASP exposure.

**Fig. 1.**
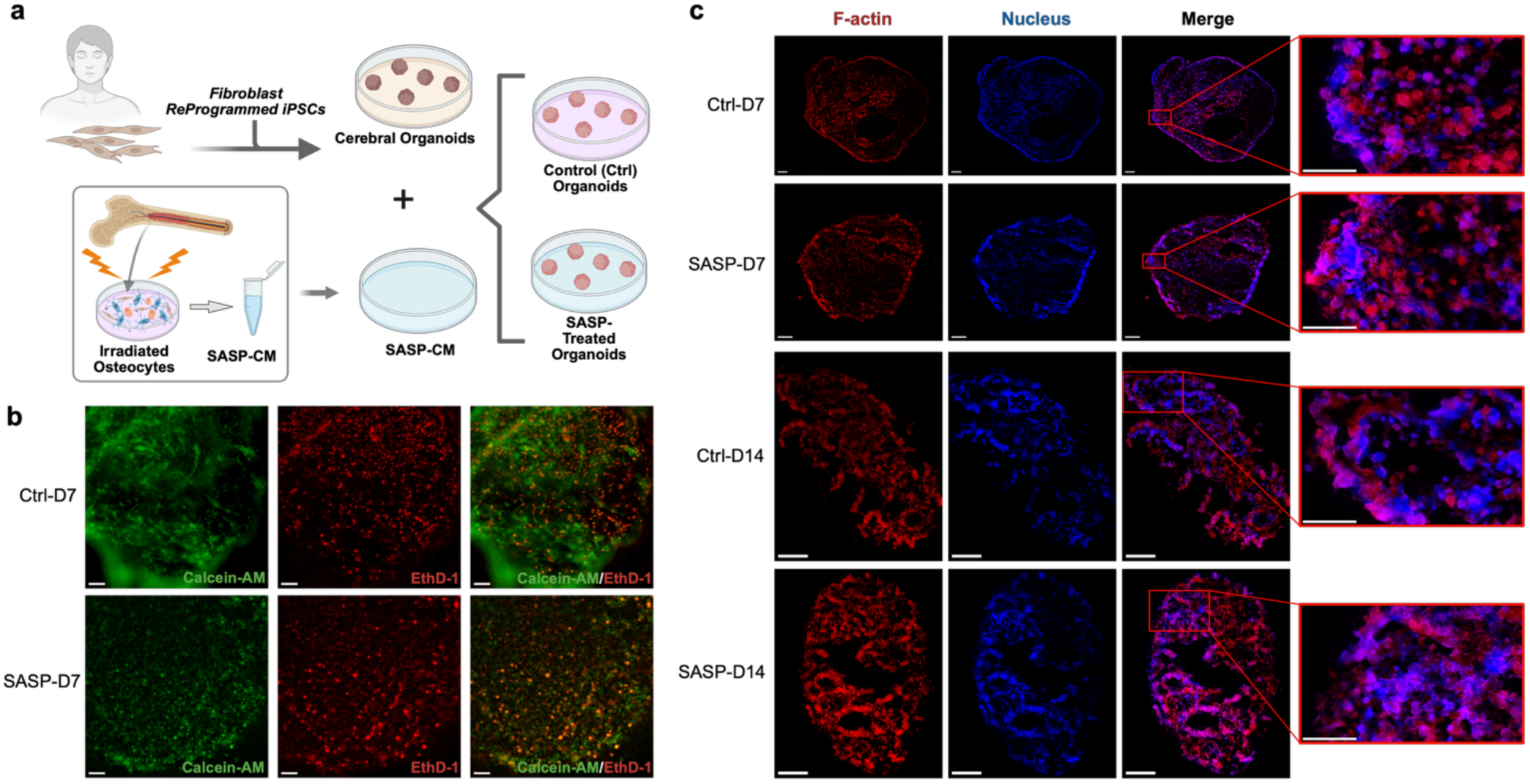
Bone-derived SASP drives progressive microarchitectural heterogeneity and F-actin/nuclear reorganization in cerebral organoids. **a**, Overview of the experimental workflow. Cerebral organoids were generated from induced pluripotent stem cells (iPSCs) reprogrammed from healthy control patient-derived fibroblasts and subsequently cultured with or without senescence-associated secretory phenotype (SASP)-enriched conditioned media. **b**, Representative Live/Dead staining images of Day 7 (D7) cerebral organoids from control and SASP-treated groups. Overview images are stitched composites and were acquired using a 10× objective lens. Scale bar: 100 µm. **c**, Representative immunofluorescence images showing F-actin (red) and nuclei (blue) across all experimental groups. Overview images are stitched composites. Images were acquired using a 20× objective lens. Scale bar: 100 µm (overview images); 50 µm (magnified regions).

**Fig. 2.**
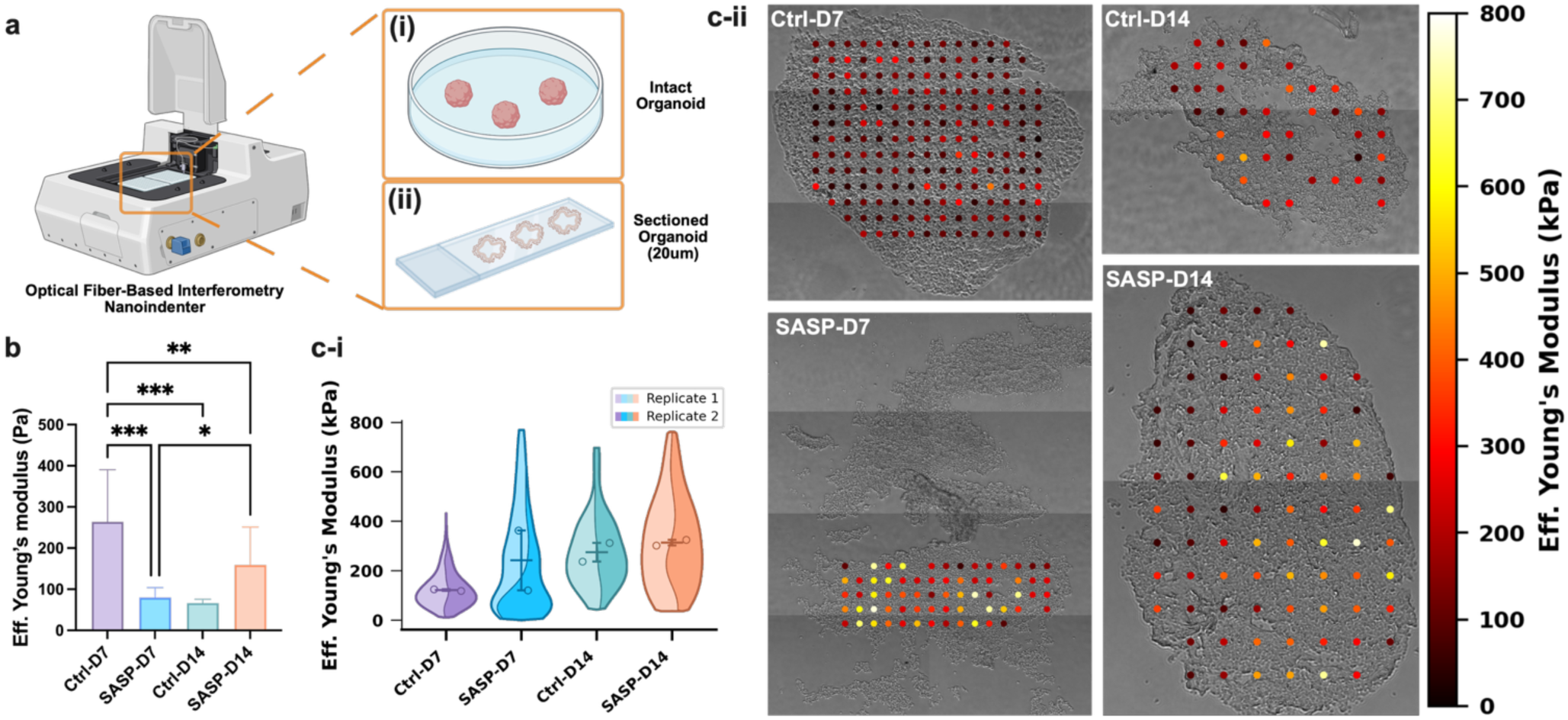
Nanoindentation-based micromechanical profiling reveals stiffness heterogeneity in live intact and cryosectioned cerebral organoids. **a**, Schematic representation of the experimental workflow for micromechanical characterization, including preparation of live and cryosectioned cerebral organoids: **i**, live intact organoids; **ii**, 20 µm cryosections. **b**, Effective elastic modulus measurements of live organoids assessed using optical fiber-based interferometry nanoindentation (N=2 biological replicates per group, 6 indentations per replicate). **c-i**, Violin SuperPlots illustrating the distribution and replicate-level statistics of effective modulus values across cryosectioned organoids (N=2 biological replicates per group, with 47–132 indentations per replicate; see Methods for details). **c-ii**, Representative high-resolution spatial stiffness maps of cryosectioned cerebral organoids. Data are presented as mean ± SD. Significance levels are denoted as *: p < 0.05, **: p < 0.01, ***: p < 0.001.

### The bone-derived SASP drives dynamic and spatially complex remodeling of cerebral organoid micromechanics

To dissect the impact of bone-derived SASP factors on the biophysical properties of human cerebral organoids, we employed two complementary micromechanical measurement approaches (Fig.2a). First, we performed live nanoindentation on intact, whole organoids under physiological (media-immersed) conditions to capture global viscoelastic responses. Second, we leveraged highly localized mapping of Young’s moduli across thin cryosectioned organoid slices (20 μm) to resolve spatial heterogeneity, including the otherwise inaccessible organoid core. All nanoindentation measurements were performed using an optical fiber-based interferometry nanoindenter (Pavone, Optics11Life). For live, intact organoids, we used a 25.5 μm radius spherical glass probe with a cantilever spring constant of 0.44 N/m and a peak load of 0.2 mN. For cryosectioned organoid samples, we employed an 11.5 μm radius spherical glass probe with a cantilever spring constant of 0.45 N/m and a peak load of 0.5 mN. These configurations allowed for sensitive, high-resolution mapping of local tissue mechanics at the cell-to-tissue scale, while minimizing mechanical perturbation to the samples^17,20,30^.

Nanoindentation of intact, live organoids (Fig.2b) revealed that control organoids experienced a significant decrease in effective Young’s modulus from D7 to D14, consistent with gradual softening during culture-associated maturation or remodeling. In contrast, SASP-exposed organoids displayed a biphasic mechanical profile; SASP-D7 exhibited significantly lower modulus than Ctrl-D7, indicating rapid early softening upon SASP exposure, but SASP-D14 underwent a significant increase in Young’s modulus relative to SASP-D7. Despite this delayed stiffening, SASP-D14 stiffness did not reach the high levels of Ctrl-D7, underscoring that chronic SASP exposure leads to an aberrant, non-physiological pattern of mechanical remodeling. These results highlight the disruptive and non-physiological trajectory of SASP-induced mechanical remodeling.

High-resolution spatial mapping on cryosectioned organoids (Fig.2c) revealed critical differences in the organization and heterogeneity of tissue mechanics. In controls, we observed a clear trend toward increased and consolidated central stiffness from D7 to D14, reflecting coordinated matrix maturation and core compaction, a hallmark of healthy organoid development. Interestingly, SASP-treated organoids at both D7 and D14 exhibited pronounced spatial heterogeneity, with numerous discrete microdomains reaching substantially higher stiffness values (up to 800 kPa) scattered throughout the tissue. This patchy and persistent distribution of hyper-stiff microdomains suggests impaired matrix remodeling and spatially restricted regions of abnormal matrix deposition in response to bone-derived SASP exposure. Rather than simply delaying or accelerating normal developmental processes, chronic SASP exposure appears to fundamentally reprogram the spatial organization of tissue mechanics, promoting the emergence and maintenance of maladaptive, fibrotic-like microenvironments^31,32^.

These findings reveal a previously unrecognized mode of SASP action: the induction of persistent, spatially complex mechanical heterogeneity within the cerebral microenvironment, reminiscent of the microenvironmental disruptions observed in the aged and diseased human brain *in vivo*^33^. Such localized stiffness “hotspots” have been implicated in impaired neural function, increased susceptibility to injury, and the propagation of degenerative cascades^34^. Our results demonstrate that bone-derived senescent secretomes can act as long-range, paracrine drivers of biophysical fragility in the brain, establishing tissue mechanics as a key mediator and potential biomarker of aging biology within the bone-brain axis.

### The bone-derived SASP triggers a pro-inflammatory molecular signature in brain organoids

To define the molecular consequences of bone-derived SASP exposure on the cerebral environment, we conducted bulk RNA sequencing of organoids harvested on Day 7. While systemic inflammation is a hallmark of aging^35,36^, the direct gene expression response of brain tissue to peripheral senescent secretomes has not, to our knowledge, been systematically characterized. We analyzed differentially expressed genes and pathway enrichments to identify key inflammatory and aging-associated transcriptional programs activated by chronic SASP exposure, providing mechanistic insight into how senescent bone cells may contribute to brain aging^37–39^. Among the samples (3 samples treated with chronic SASP exposure and 3 control samples untreated), 30,605 genes were detected. 2477 differentially expressed genes (DEGs) were identified, of which 1700 were upregulated and 777 were downregulated.

Unsupervised clustering and functional enrichment analyses revealed that SASP exposure elicited broad upregulation of gene sets associated with ossification, endochondral bone development, actin filament bundling, and interleukin signaling, suggesting that bone-secreted factors may activate morphogenic and immune-related pathways in brain organoids (Fig.3a). Additional enrichment was observed for terms linked to wound healing, substrate adhesion, and vascular remodeling, indicating broad activation of tissue remodeling and structural adaptation programs. In contrast, downregulated genes were strongly enriched for processes involved in neurodevelopment, synaptic signaling, and neurotransmitter secretion. Major enriched categories included glutamatergic synapse function, dopamine secretion, and vesicle-mediated transport, indicating suppression of neuronal communication machinery. Furthermore, clusters associated with RNA polymerase binding, neuronal cell body organization, and associative learning and memory suggest a broad decline in transcriptional and cognitive gene programs. The suppression of glutamate receptor signaling and synaptic vesicle cycle genes points toward impaired synaptic plasticity and neurotransmission in the SASP-treated organoids.

Hierarchical clustering further confirmed consistent and reproducible transcriptomic shifts across biological replicates (Fig.3b). Notably, several genes upregulated in SASP-treated brain organoids, including IGFBP3, B2M, and SERPINE1 are secreted factors or ECM modulators previously implicated in aging-related neuroinflammation and cognitive dysfunction^40–42^. Conversely, key neural developmental genes such as MYT1_1 and MARF1 were downregulated, potentially indicating altered neurodevelopmental trajectories in response to bone-derived secretions^43,44^. These findings suggest a mechanistic link between the bone secretome and molecular pathways relevant to brain aging and degeneration.

Gene ontology (GO) and KEGG pathway enrichment analysis of differentially expressed genes revealed distinct biological programs modulated by chronic SASP exposure in brain organoids (Fig.4). Among upregulated genes, enrichment terms were dominated by those related to extracellular matrix (ECM) organization, wound healing, cell adhesion, and connective tissue development. These findings suggest activation of structural remodeling pathways in the treated organoids. Terms such as “collagen-containing extracellular matrix,” “cell-substrate adhesion,” and “focal adhesion” indicate increased engagement with matrix remodeling and mechanotransduction processes^45–50^. This was further supported by KEGG pathway enrichment of PI3K-Akt signaling, ECM-receptor interaction, and AGE-RAGE signaling in diabetic complications, all of which are pathways known to mediate inflammatory remodeling, tissue integrity, and age-related cellular stress responses^50–55^.

**Fig. 3.**
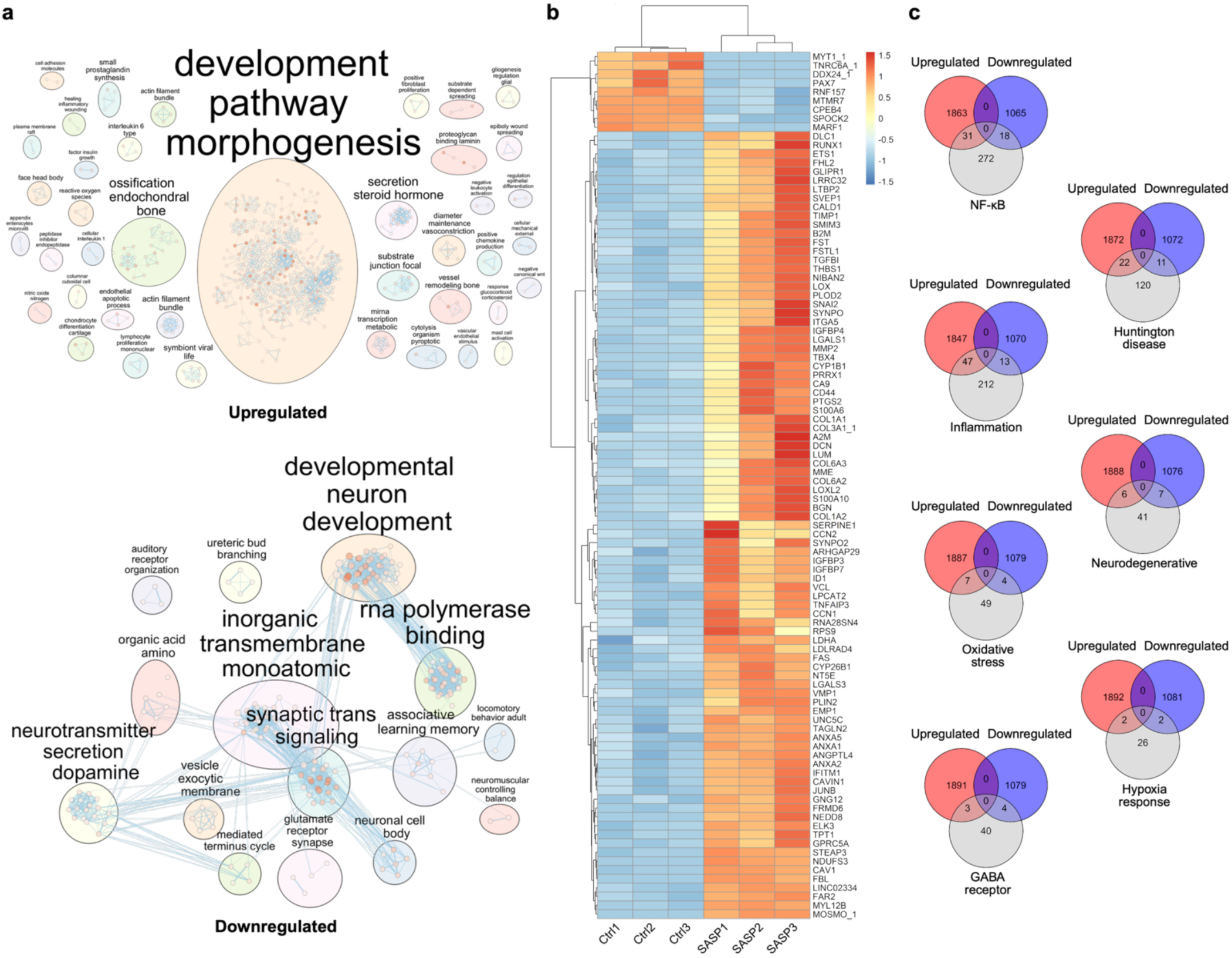
ECM and inflammatory gene upregulation, synaptic pathway repression, and stress signature enrichment by bulk RNA-seq. **a**, Pathway enrichment analysis of the SASP organoids separated by upregulated genes (top) and downregulated genes (bottom). **b**, Sample heatmap and dendrogram of the most significantly differentially expressed genes based on the adjusted p-value. **c**, Venn diagrams comparing numbers of differentially expressed genes between SASP upregulated genes and downregulated genes and differentially expressed genes from Allen Brain Atlas microarray data for collection of classifications and diseases. N=3 biological replicates, 3 treated with chronic SASP exposure and 3 samples untreated (controls).

**Fig. 4.**
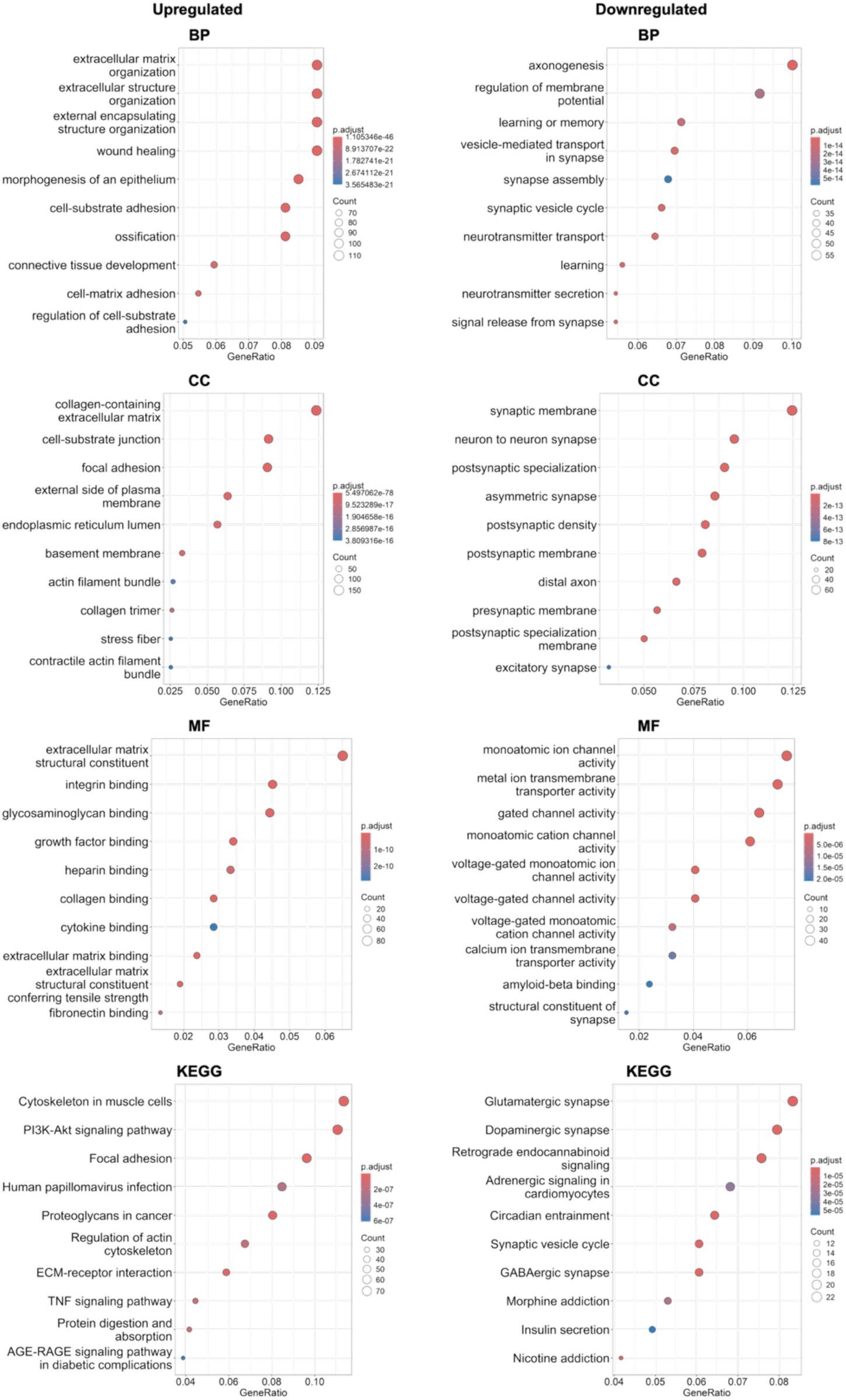
SASP exposure alters gene programs in brain organoids revealed by GO and KEGG enrichment analysis. Scatter plots of the top ten GO terms for biological pathways (BP), molecular function (MF) and cellular components (CC) as well as KEGG pathways, separated by upregulated (left) and downregulated (right) genes. N=3 biological replicates, 3 treated with chronic SASP exposure and 3 samples untreated (controls).

Conversely, the downregulated gene set was significantly enriched for terms involved in synaptic signaling, neurotransmitter secretion, learning and memory, and axonogenesis. The suppression of these terms suggests an impairment in neuronal communication and plasticity^56–59^. GO terms like “synaptic vesicle cycle,” “neurotransmitter transport,” and “regulation of membrane potential” were among the most enriched, pointing to broad dysfunction in signal transmission machinery. KEGG analysis further revealed the downregulation of key neuronal circuits including the glutamatergic, dopaminergic, and GABAergic synapse pathways, as well as circadian entrainment and insulin signaling. These pathways are central to cognitive performance, emotional regulation, and metabolic-brain interactions, systems often disrupted during brain aging and neurodegeneration^60–66^.

To evaluate the relevance of our DEGs to established brain aging and disease-related pathways, we performed gene set intersection analysis using microarray datasets from the Allen Brain Atlas (Fig.3c)^67,68^. Venn diagrams revealed substantial overlap between our SASP-induced DEGs and curated gene sets related to NF-κB signaling, inflammation, and oxidative stress, with 31, 47, and 7 upregulated genes intersecting, respectively. These categories are known mediators of neuroinflammation and cellular senescence^67–69^, suggesting that the bone secretome activates conserved aging-related molecular programs in the brain. Additionally, upregulated DEGs overlapped with genes associated with Huntington’s disease^70^, neurodegenerative pathways^67,71^, and hypoxia response^72,73^, indicating potential involvement of broad neurodegenerative mechanisms. Furthermore, there was intersection between our downregulated DEGs and gene sets related to GABA receptor signaling, emphasizing potential disruptions in inhibitory neurotransmission. Collectively, these results indicate that SASP exposure to brain organoids recapitulates transcriptional changes linked to both inflammatory activation and neuronal dysfunction seen with aging and disease.

The upregulation of ECM, collagen synthesis, and TNF/inflammatory pathways mechanistically underpins the biphasic mechanical remodeling and emergence of spatial heterogeneity observed in our organoid model. Local increases in ECM gene expression, combined with sustained inflammatory signaling, likely drive both the heterogeneous stiffening and cytoskeletal disorganization observed in the data in Fig.1 and 2. Conversely, the downregulation of synaptic and neurotransmitter-related pathways may predispose the neural network to reduced plasticity and increased vulnerability to further stressors, mirroring early events in age-associated brain decline. Taken together, these results provide molecular evidence that bone-derived senescent secretomes can reprogram the human brain microenvironment in a paracrine fashion by engaging conserved inflammatory, stress, and neurodevelopmental vulnerability networks.

### Activation of pro-inflammatory and senescence-associated markers in cerebral organoids

To validate and spatially localize the pro-inflammatory and senescence responses predicted by our transcriptomic analysis, we performed multiplex immunofluorescence for key pathway effectors in cerebral organoids following seven days of exposure to bone-derived SASP compared to control condition (Fig.5). Immunofluorescence imaging (Fig. 5a, b) revealed changes in the spatial distribution of IL-1β following SASP exposure, with staining appearing more broadly distributed across the organoid area in the SASP-D7 group compared to Ctrl-D7. However, quantitative analysis of the fluorescent area did not show a statistically significant difference between SASP-D7 and Ctrl-D7 (Fig. 5c), likely reflecting biological variability among organoids and heterogeneous responses to SASP. Similarly, IL-6 did not exhibit a significant change in response to SASP treatment. These results suggest that while SASP exposure may influence the spatial pattern of certain inflammatory markers, it does not lead to a measurable overall change in IL-1β or IL-6 signal intensity within the 7-day timeframe.

**Fig. 5.**
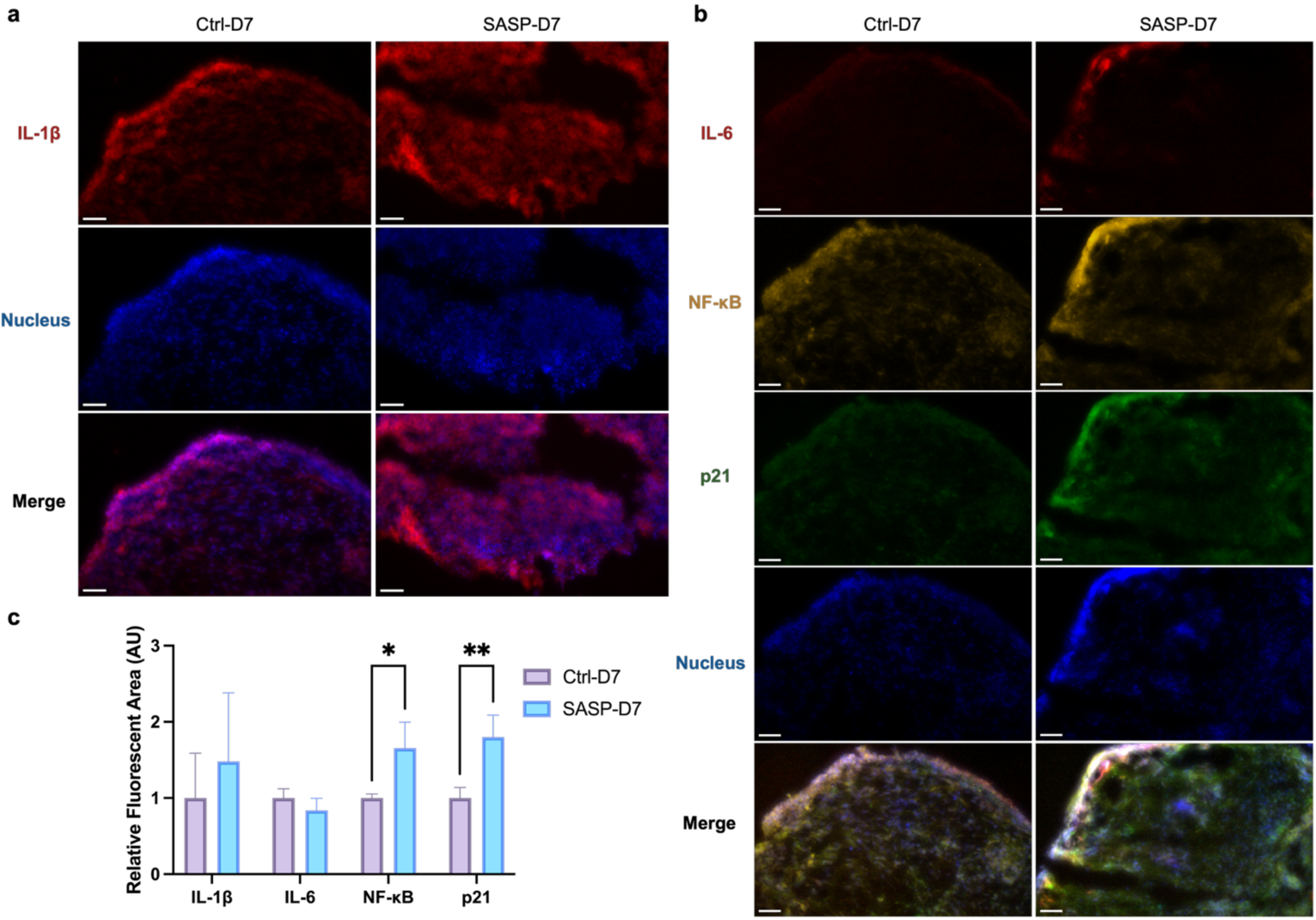
Immunofluorescence analysis of cerebral organoids following SASP-CM treatment. **a**, Representative immunofluorescence staining of cerebral organoids showing IL-1β (red) and nuclei (blue, DAPI). **b**, Representative multiplex immunofluorescence staining of cerebral organoids showing IL-6 (far-red), NF-κB (yellow), p21 (green), and nuclei (blue, DAPI). **c**, Quantification of fluorescence area for IL-1β, IL-6, NF-κB, and p21. Fluorescent regions were measured using Fiji. Values were normalized to the mean area of the Ctrl-D7 group for each marker. For each group and marker, N=4-5 micrographs were analyzed per condition and time point. Data are shown as mean ± SD. Significance levels are denoted as *: p < 0.05, **: p < 0.01. All Images in the figure were acquired using a 20× objective lens with a scale bar: 50µm.

In contrast to IL-1β and IL-6, NF-κB fluorescence (*p*< 0.05), indicating strong activation of inflammatory transcriptional programs. As a master regulator of the SASP, NF-κB orchestrates the expression of a wide array of pro-inflammatory mediators and senescence-associated genes^74^. The significant upregulation of NF-κB, even in the absence of significant changes in IL-1β and IL-6, highlights that SASP exposure robustly triggers transcriptional inflammatory pathways independently of proportional changes in upstream cytokine abundance. This observation aligns with models where NF-κB activation serves as an early and sustained driver of senescence-associated inflammation, potentially maintained through autocrine and paracrine feedback mechanisms^75,76^. The data suggest that SASP exerts its pro-inflammatory influence primarily through NF-κB-mediated transcriptional activation rather than direct amplification of soluble cytokines like IL-6.

Bone-derived SASP treatment also significantly increased p21 expression, a core marker of cellular senescence, supporting the induction or reinforcement of a senescence-like program within the organoid microenvironment. The coordinated elevation of both pro-inflammatory and senescence markers is notable, as this suggests not only acute paracrine reprogramming but also the possibility of a feed-forward loop where SASP factors induce secondary senescence and propagate inflammatory stress among neural and non-neural cells alike^77–79^. This mechanism has been implicated in the spread of tissue dysfunction in both aging and neurodegenerative contexts^77,80^, and our data extend this concept to the brain-bone signaling axis.

In contrast to D7, the D14 group showed statistically significant increases in all four markers, including IL-1β, IL-6, NF-κB, and p21, following SASP exposure (Extended Data Fig.3). Immunofluorescence imaging revealed markedly enhanced signaling and broader spatial distribution in SASP-treated D14 organoids compared to controls. Quantitative analysis confirmed significant elevations in both pro-inflammatory cytokines and senescence-associated markers. These results suggest that the inflammatory and senescence responses become more pronounced with prolonged paracrine SASP exposure, indicating a time-dependent amplification of SASP-induced effects in cerebral organoids. The coordinated elevation of these key markers, together with strong transcriptomic evidence for inflammatory and stress pathway engagement, underscores the robustness of the SASP-driven phenotype.

Altogether, our findings position the bone-derived SASP factors as a potent extrinsic driver of neuroinflammatory and senescence responses in the human brain, mechanistically linking musculoskeletal aging to neural vulnerability. To our knowledge, this is the first study to directly demonstrate that bone-derived senescent secretomes can remodel the neuroinflammatory and mechanical microenvironment of human cerebral organoids via paracrine signaling. We acknowledge that our study design involved the use of murine-derived senescent osteocyte secretome to treat human cerebral organoids. This approach is consistent with long-standing practices in cell biology, where animal-derived sera (e.g., fetal bovine serum) and matrices (e.g., Matrigel) are routinely used to support human cell and organoid culture^81–83^. Importantly, the key molecular effectors of the SASP, including IL-6, IL-1β, and other cytokines, are highly conserved across mammals, supporting the validity of our mechanistic insights^84^. Nonetheless, species-specific differences in secretome composition or receptor-ligand interactions could modulate some responses. Addressing this limitation, our future studies will leverage human-derived primary senescent bone cells or fully humanized co-culture systems to confirm and extend the translational relevance of our platform.

## Implications and Future Directions

Our study provides evidence that bone-derived senescent secretomes can reprogram the neural microenvironment in a paracrine manner, inducing robust neuroinflammatory, mechanical, and molecular aging signatures in human cerebral organoids. This mechanistic insight advances the emerging paradigm that musculoskeletal aging can act as a long-range, extrinsic driver of brain vulnerability, a concept recently suggested by preclinical studies but not to our knowledge previously demonstrated using human cross-organ models^85,86^. Unlike prior work in animal models or plasma parabiosis studies^86,87^, our tractable organoid platform enables direct interrogation of human-specific bone**-**brain interactions and dissection of causal pathways.

Building on these findings, a critical next step is to deconvolute the SASP to identify the specific bioactive factors (i.e., cytokines, proteases, chemokines, and extracellular vesicles) most responsible for reprogramming brain tissue mechanics and the inflammatory state^88,89^. While canonical factors like IL-1β and IL-6 are clearly implicated^90,91^, the complexity of the senescent secretome suggests combinatorial or synergistic effects that remain to be resolved. Leveraging single-cell transcriptomics, spatial proteomics, and high-content longitudinal imaging will be essential for delineating which neural cell types or developmental stages are most susceptible to peripheral senescence cues and for mapping cell-specific responses that underlie tissue-level fragility. Integration of vascular, blood**-**brain barrier, or immune elements into future multi-organ platforms will further refine our understanding of how systemic aging signals propagate to the brain and may reveal critical checkpoints for therapeutic intervention^87,92^. Recognizing the inherent limitations of our current reductionist system, our ongoing work is focused on advanced 3D bioprinted cerebral microenvironments. By spatially organizing organoids within engineered, perfusable matrices and enabling precise, tunable delivery of bone-derived SASP factors, this platform will allow us to model physiological diffusion gradients, dissect spatiotemporal exposure dynamics, and more accurately recapitulate the complexities of *in vivo* bone-brain signaling. These advances will enable direct investigation of how systemic skeletal aging signals shape neural microenvironments at single-cell and spatially resolved levels.

Clinically, these advances underscore musculoskeletal health, in particular bone senescence, as modifiable, upstream determinants of brain resilience. This opens new translational avenues: targeting peripheral senescence, through senolytic agents, SASP-modifying drugs, or bone**-**brain axis interventions, could represent a paradigm shift in the prevention or treatment of cognitive decline and neurodegeneration. More broadly, our work establishes a new platform for testing cross-organ aging hypotheses in human systems, with a potential impact extending well beyond the bone**-**brain axis to other age-related diseases with systemic drivers.

## Experiments

### Preparation and Validation of Senescent Osteocyte-Conditioned Medium (SASP-CM)

Senescent osteocyte-conditioned medium (SASP-CM) was generated using MLO-Y4 osteocyte cultures, as previously described and validated in our recent published study^20^. Briefly, immortalized mouse-derived osteocytes (MLO-Y4, sourced from the Professor Linda Bonewald laboratory) were used to ensure reproducibility and to minimize variability inherent to heterogeneous primary bone cell cultures. Cells were cultured on rat tail collagen type I-coated plates and passaged twice prior to senescence induction^17,20^. Senescence was induced *in vitro* by a single 10 Gy dose of X-ray irradiation (CellRad, Precision X-Ray, Inc.), a method chosen for its well-established reproducibility and ability to reliably trigger DNA damage response, leading to stable cell cycle arrest. For the preparation of conditioned medium, irradiated osteocyte cultures were maintained for 20 days post-irradiation to allow for full establishment of the senescent phenotype (Extended Data Fig. 1)^17,20^. On day 20, the culture medium was collected, centrifuged at 1500 x g for 5 min to remove cellular debris, aliquoted, and stored at −80 ℃ until use. The senescent state was confirmed by nearly 100% SA-β-gal positivity and significant upregulation of SASP factors, including *Il-6*, *Tnf-α*, and *Mmp12* (Extended Data Fig. 1). The same batch of SASP-CM was used in all experiments presented in this study.

### Cerebral Organoid Generation and Culture

Control healthy human iPSC Cells obtained from (NIMH Repository & Genomics Resource Cell line MH0185984) were expanded in mTESR plus media (StemCell Tech) on vitronectin-coated surfaces. Initial cell expansion occurred in T25 culture vessels before scaling up to T75 flasks for routine maintenance. Subculturing was performed every 5-7 days using a 1:2 split ratio. The passaging procedure involved treating confluent cultures with Versene (Gibco) for 4 minutes at 37°C, followed by gentle mechanical dissociation using cell scrapers to lift the colonies. To ensure genomic stability and maintain pluripotency characteristics, all iPSC lines were kept under passage number 50 throughout the experimental period. Quality control measures included routine mycoplasma contamination screening, with all cell lines confirmed negative before use in downstream applications.

Human cerebral organoids were derived from control induced pluripotent stem cells (iPSCs) following a 14-day differentiation protocol with daily medium exchanges. Initially, iPSC colonies were prepared for dissociation by washing with 1X Hanks’ Balanced Salt Solution (HBSS, Gibco), then enzymatically dissociated into single-cell suspensions using Accutase treatment. For embryoid body (EB) formation, dissociated cells were plated at a density of 25,000 cells per well in ultra-low attachment 24-well Aggrewell plates (800mm, Stem Cell Technologies). During the initial 24-hour period, EBs were cultured in Essential 8 medium containing DMEM/F12, L-glutamine, and sodium bicarbonate (Gibco), with the addition of Y-27632 ROCK inhibitor (10µM, 1:1000 dilution) to enhance cell survival. Neural fate specification was initiated by transitioning cultures to a neural induction medium supplemented with dual SMAD pathway inhibitors. This medium consisted of Neural Basal Medium with N2 and B27 supplements, containing 10µM SB431542 (TGFβ pathway inhibitor) and 0.1µM LDN193189 (BMP pathway inhibitor). EBs remained under these neural-inducing conditions for 5-7 days to drive neuroectoderm commitment. Subsequently, neural progenitor expansion was promoted by switching to neural differentiation medium, which maintained the Neural Basal Medium base with N2 and B27 supplementation but excluded the SMAD inhibitors. This phase lasted 4-5 days to allow robust neural progenitor proliferation. On day 14, three-dimensional architectural support was provided by embedding the developing neural spheroids in Matrigel droplets. Following embedding, organoids were transferred to specialized cerebral organoid maturation medium (Catalog # 08571) designed to support diverse neural cell type development and the establishment of organized neural tissue architecture. All assays were performed at 60 days of differentiation; this is when we see electrical activity emerging. All cell types are also seen at day 60, confirmed via bulk and single nuclear RNA seq in previous publication^93^.

### Treatment of cerebral organoids

Following initial maturation, the organoids were transferred from the Kathuria Lab (Johns Hopkins University) to the Tilton Lab (University of Texas at Austin) on the same day. Upon receipt, organoids were immediately transferred to well plates and allowed to recover for 48 hours in the same cerebral organoid maturation medium used prior to shipping. After the 48-hour recovery phase, organoids were randomly divided into two groups: (1) a control group, for which the medium was changed to α-Minimum Essential Medium Eagle (αMEM, Sigma-Aldrich) supplemented with 5% fetal bovine serum (FBS, Gibco), 5% calf serum (CS, Sigma-Aldrich), and 1% penicillin–streptomycin (10,000 U/mL, Gibco); and (2) a SASP-CM group, for which the medium was changed to osteocyte-derived SASP-conditioned medium prepared in the same complete αMEM background media. Media changes were performed every 2-3 days throughout the treatment period.

Organoids from both study groups were collected on days 7 and 14 post-treatment for downstream analyses. For cryosectioning, organoids were washed and fixed in 4% paraformaldehyde (PFA) for 1 hour at room temperature, followed by three washes with Dulbecco’s phosphate-buffered saline (DPBS; Gibco). Samples were then embedded in an OCT compound and cryosectioned using a CryoStar NX70 cryostat microtome (Epredia, Fisher Scientific). Sections were cut at 10 µm thickness for immunostaining and 20 µm thickness for micromechanical profiling. In parallel, intact live organoids from each group and time point were subjected to non-destructive nanoindentation assays. Additionally, three biological replicates per group at each time point were collected for RNA analysis. For RNA extraction, organoids were homogenized in TRIzol reagent (Invitrogen), and total RNA was purified using the GeneJET RNA Purification Kit (Thermo Fisher Scientific) according to the manufacturer’s protocols. RNA concentrations were measured using a NanoDrop Microvolume Spectrophotometer (Thermo Fisher Scientific), and purified RNA was processed for downstream bulk RNA sequencing.

### Assessment of organoid viability following SASP exposure

Organoids were washed three times with sterile DPBS (Gibco) before staining. A working dye solution was prepared using the Live/Dead Viability/Cytotoxicity Kit (Invitrogen) by adding 2 µL of 2 mM Ethidium Homodimer-1 and 0.5 µL of 4 mM calcein AM to 5 mL of DPBS, yielding final concentrations of 0.8 µM and 0.4 µM, respectively. The solution was vortexed and applied to the organoids, which were then incubated at room temperature for 45 minutes in the dark. After incubation, samples were washed once with DPBS prior to imaging.

### Assessment of actin cytoskeletal organization following SASP exposure

Cryosectioned organoid slides (10 μm) were thawed at room temperature for 15 minutes prior to staining. A hydrophobic barrier was drawn around each sample using the Aqua-Hold Pap Pen 2 (Fisher Scientific) to confine reagents during staining. Samples were washed twice with pre-warmed PBS to remove residual OCT and equilibrate the sections. For viability assessment, CellMask Deep Red Actin Tracking Stain (Thermo Fisher Scientific) was diluted in PBS (1:500 dilution). The staining solution was applied to the samples and incubated for 1 hour at room temperature. The nuclei were counterstained with 4′,6-diamidino-2-phenylindole (DAPI, Invitrogen) at 1X concentration for 10 minutes. Slides were then washed with PBS prior to imaging.

### Biomechanical remodeling of cerebral organoids following SASP exposure

Nanoindentation testing was performed on both intact (live) cerebral organoids and 20 µm-thick cryosectioned sections using an optical fiber-based interferometry nanoindenter (Pavone, Optics11 Life). For live intact organoids, a probe with a 25.5 µm radius and 0.44 N/m stiffness was used; indentations were conducted in Peak Load Poking (PLP) mode with a maximum force of 0.2 µN. For sectioned samples, an 11.5 µm radius probe (0.45 N/m stiffness) was used. Indentations were performed across a 50 µm × 50 µm grid, with a peak load of 0.5 µN and a piezo speed of 30 µm/s. The apparent effective Young’s modulus for all sample types was calculated using Hertzian contact mechanics to assess local tissue mechanical properties. Indentation force-displacement curves were analyzed using DataViewer V2.5.2 (Optics11 Life), with curve fitting restricted to the linear loading phase for accurate modulus extraction.

#### Statistical analysis

All statistical analyses were performed in GraphPad Prism (v10) and Python (superviolin package)^94^. Separate analyses were conducted for (a) apparent effective Young’s modulus values from intact live organoid measurements and (b) spatial mapping of modulus across cryosectioned organoid grids.

##### (a) Intact organoid nanoindentation

Statistical differences among the study groups (i.e., Ctrl-D7, SASP-D7, Ctrl-D14, and SASP-D14) were assessed by one-way analysis of variance (ANOVA) followed by Tukey’s post hoc multiple comparisons test. Prior to ANOVA, assumptions of homogeneity of variance were tested using both Brown-Forsythe and Bartlett’s tests; both indicated significant differences in variance across groups (P < 0.001 for each). Statistical significance was defined as p < 0.05. The ANOVA test indicated a significant main effect of group, with post hoc Tukey’s tests showing significant differences between Ctrl-D7 and SASP-D7 (p < 0.001), Ctrl-D7 and Ctrl-D14 (p < 0.001), Ctrl-D7 and SASP-D14 (p = 0.009), and SASP-D7 and SASP-D14 (p = 0.030).

##### (b) Cryosectioned grid spatial mapping

To visualize spatial and replicate-level heterogeneity, effective modulus values from each indentation coordinate within the 50 µm x 50 µm grid were analyzed using the superviolin Python package^94^, which overlays replicate means and error bars on violin plots for biological replicates. Kernel density estimation (KDE) smoothing bandwidth was set by Scott’s factor, with fitted values ranging 0.414-0.608 (within the recommended 0.4-0.6 interval), indicating appropriate smoothing. Subsequently, one-way ANOVA was used to compare modulus distributions across groups (P = 0.304, n.s.), followed by post hoc Tukey’s HSD tests (all pairwise p > 0.28).

### Immunofluorescence analysis of structural and inflammatory markers

Tissue cryosections were outlined with a hydrophobic PAP pen (Aqua-Hold, Fisher Scientific) to contain reagents during staining. Slides were rehydrated in DPBS and incubated for 1 hour at 37 °C in a humidified, dark slide box with a blocking buffer consisting of 5% bovine serum albumin (BSA) in DPBS. For multiplex staining, primary antibodies were diluted 1:200 in antibody solution (5% BSA, 0.5% Triton X-100 in DPBS) and incubated overnight at 4 °C. The primary antibody mixture included mouse IgG1 anti-p21 (WA-1), mouse IgG2a anti-IL-6 (OTI3G9), and rabbit anti-NF-κB. Following incubation, slides were rinsed with DPBS and washed for 3 × 10 minutes in PBST (DPBS with 0.1% Tween 20) on an orbital shaker at room temperature. Secondary antibodies were diluted 1:1000 in antibody solution and applied for 2 hours at room temperature. For multiplex detection, Alexa Fluor 488 goat anti-mouse IgG1 (p21), Alexa Fluor 647 goat anti-mouse IgG2a (IL-6), and Alexa Fluor 555 goat anti-rabbit IgG F(ab’)₂ (NF-κB) were used. DAPI (2 μg/mL in PBS) was applied for 10 minutes for nuclear staining, followed by the same wash procedure and a final rinse in PBS. Slides were mounted with antifade medium, cured overnight, and stored at 4 °C until imaging. IL-1 staining was performed separately using the same protocol. Sections were incubated with rabbit anti-IL-1 (1:200), followed by Alexa Fluor 555 goat anti-rabbit IgG F(ab’)₂ (1:1000) under identical conditions. Images were acquired using a 20x objective lens under identical acquisition settings across conditions. For quantification, background-subtracted images were analyzed by placing a region of interest (ROI) over each images to quantify the integrated density using Fiji^95–97^.

#### Statistical analysis

For each marker, three biological replicates were analyzed per experimental group and time point. Raw area measurements from SASP-D7 were normalized to the mean area of the Ctrl-D7 group for each marker. Normalized values were used to compute group means and standard deviations (SD). Statistical analyses were performed in GraphPad Prism. Normalized fluorescence-area values for all four markers were analyzed by two-way ANOVA. The main effects for treatment and marker were interpreted, and marker-specific differences were identified by Tukey’s multiple-comparisons test. All tests were two-tailed, with significance set at p< 0.05. The same statistical analysis workflow described above for the D7 time point was applied to the D14 experimental groups (Ctrl-D14 and SASP-D14) (Extended Data Fig. 3).

### Transcriptomic profiling of SASP-induced changes in cerebral organoids

#### RNA Preparation

RNA integrity was assessed using an Agilent 4200 Tapestation, ensuring RNA Integrity Number (RIN) values above 5.0 for all samples. A total of 150 ng of RNA per sample was used as input for library preparation.

#### Library Preparation

Libraries were prepared using the NEBNext UltraExpress RNA Library Prep Kit (New England Biolabs, Ipswich, MA) following the manufacturer’s protocol. mRNA was enriched using poly(A) selection. Enriched mRNA was fragmented, reverse-transcribed, and converted to cDNA. cDNA underwent end repair, adapter ligation, and size selection. PCR amplification was performed using 11 cycles to ensure sufficient library yield while minimizing amplification bias. Library quality and size distribution were assessed using a Tapestation (Agilent) and quantified by Qubit (Thermo Fisher Scientific).

#### Sequencing

Final libraries were pooled equimolarly and sequenced on an Illumina NovaSeq X Plus platform using a 25B lane configuration with paired-end 150 bp (PE150) reads with a minimum of 20M Reads per sample. The sequencing run achieved an average depth of 20M reads per sample. Base calling and demultiplexing were performed using Illumina’s DRAGEN Bio-IT platform.

#### Bulk RNA-Seq Analysis

Raw RNA-seq data were processed using a standard pipeline that included quality control, trimming, alignment, and transcript quantification. Sequencing quality was assessed with FastQC^98^, and summary reports were compiled using MultiQC^99^. Adapter sequences and low-quality bases were trimmed using Trim Galore^100^. Reads were aligned to the reference genome using STAR^101^, and transcript-level quantification was performed using Salmon^102^ in alignment-based mode, generating quant.sf files for each sample. These quantifications were subsequently imported into R using the tximport^103^ package for gene-level summarization prior to differential expression analysis.

#### Differential Gene Expression Analysis

Differentially expressed genes (DEGs) were identified using the DESeq2 R package (version 1.46.0)^104^. The dataset was filtered by a cutoff of adjusted *p*-value < 0.05 and absolute fold change > 2. DEGs were subjected to GO enrichment analysis and KEGG pathway analysis using the clusterProfiler R package (version 4.16.6)^105^. The enrichment network map was created by feeding DEGs into g:Profiler^106^ and retrieving GO terms size 5-200. The enrichment network map was visualized in Cytoscape (version 3.10.3)^107^ using the EnrichmentMap^108^ and AutoAnnotate^108^ modules. Venn diagrams were created by downloading microarray data from Allen Brain Atlas for classifications and diseases of interest and comparing their DEGs to our DEGs.

## Acknowledgements

Figure schematics and assemblies of quantified and analyzed data were done with BioRender.com license to BioMATTER Lab (Tilton group). Bulk RNA sequencing was performed by 7-Traits Genomics through a service contract. The authors also acknowledge supports from the core facilities at The University of Texas at Austin and Johns Hopkins University.

## Author Contributions

M.T. and A.K. developed the concept and designed the research project. C.K. and M.T. performed the bone cell senescence, mechanobiological experiments, SASP exposure studies, and interpreted the results. L.D.O., A.Ks., and S.K. generated cerebral organoids to maturation at the Kathuria Lab (JHU), which were then used in all the downstream bone-derived SASP experiments by C.K. at the BioMATTER Lab (UT Austin). L.D.O. analyzed bulk RNA sequencing data. C.K., M.T., and A.Y.L. designed the optical fiber interferometry nanoindentation experiments. C.K. performed and analyzed nanoindentation experiments and immunofluorescent assays. C.K. and M.T. wrote the original manuscript; L.D.O. and A.K. wrote sections on organoid generation and bulk RNA sequencing. J.L.K. provided critical intellectual input and contributed to funding acquisition. All authors read, edited, and approved the final version of the manuscript.

## Data Availability Statement

The data that support the findings of this study are available on request from the corresponding authors.

