## Supplemental Materials for "Paracrine Bone-Derived Senescent Secretome Induces Spatially Patterned ECM and Biomechanical Vulnerability in Human Brain Organoids"

*Corresponding authors:

### Extended Data

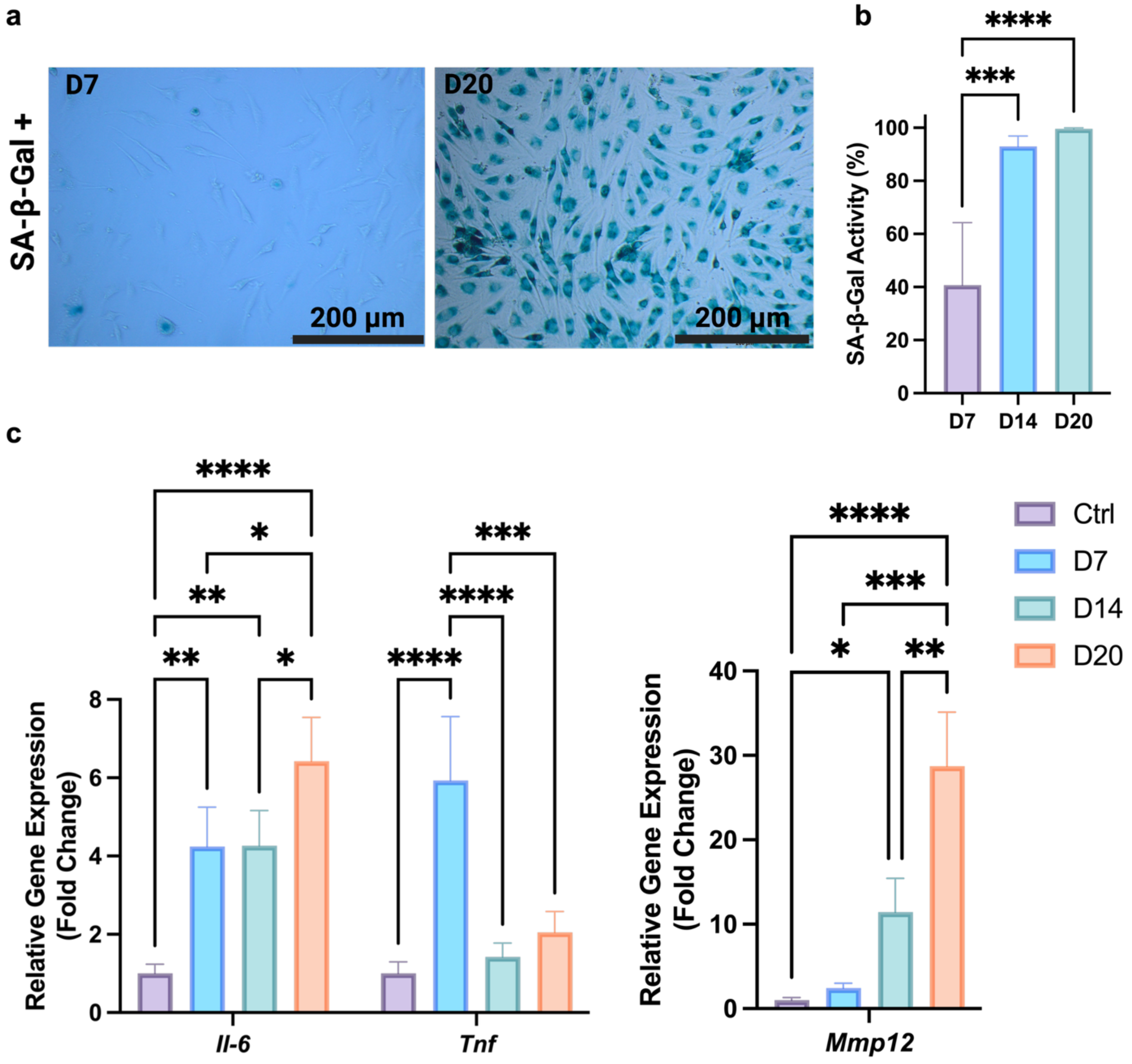


**Extended Data Fig. 1| Validation of senescent phenotype and SASP production in irradiated osteocyte cultures used for SASP-CM generation. a,** Representative images of SA-β-Gal staining osteocytes cultures post-irradiation on days 7 and 20. **b,** Quantification of SA-β-Gal + cells as a percentage of total cells. **c,** RT-qPCR analysis of SASP gene expression (*Il-6, Tnf-α, Mmp12*) in irradiated versus control non-senescent cultures on day 20, normalized to Gapdh (Housekeeping gene) and shown as fold change relative to control. Data are presented as mean ± SD for N=3 biological replicates. Significance levels are denoted as *: $p < 0.05$, **: $p < 0.01$, ***: $p < 0.001$. ****: $p < 0.0001$. **All analyses were performed on the same batch of irradiated cultures used for SASP-CM generation in this work.** Panels **a-c** are reproduced from Tilton et al., *Small* 2025, with permission, to validate the preparation of SASP-CM used in the present study.

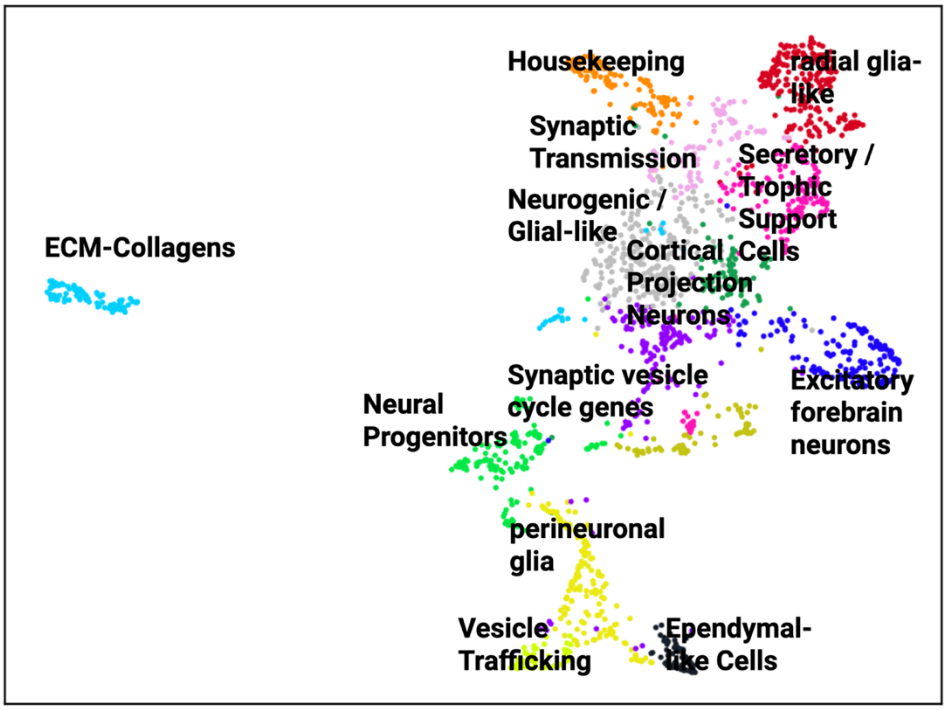


**Extended Data Fig. 2| Single-nucleus transcriptomic profiling of 2-month-old cerebral organoids reveals diverse neuronal and non-neuronal populations.** UMAP visualization of single-nucleus RNA-seq data from 2-month-old cerebral organoids identifies 14 transcriptionally distinct clusters, each annotated by enriched gene signatures. Annotated populations include excitatory forebrain neurons (e.g., *GRIA2*$^{+}$, *RBFOX3*$^{+}$), synaptic vesicle cycle-enriched neurons (*SNAP25*$^{+}$, *PCLO*$^{+}$), cortical projection neurons (*NRXN1*$^{+}$, *TBR1*$^{+}$), and multiple non-neuronal subtypes such as neurogenic/glial-like cells (*NEAT1*$^{+}$, *RMST*$^{+}$), radial glia-like progenitors (*MIAT*$^{+}$, *MBD5*$^{+}$), and vesicle trafficking-associated trophic support cells (*VEGFA*$^{+}$, *TRPM7*$^{+}$). ECM clusters (*COL1A1*$^{+}$, *PRRX1*$^{+}$) and ECM-producing cells are also detected, alongside a small population of motile ciliated (ependymal-like) cells. This transcriptional landscape demonstrates advanced neuronal subtype emergence and, characteristic of developing cerebral organoid at this stage.

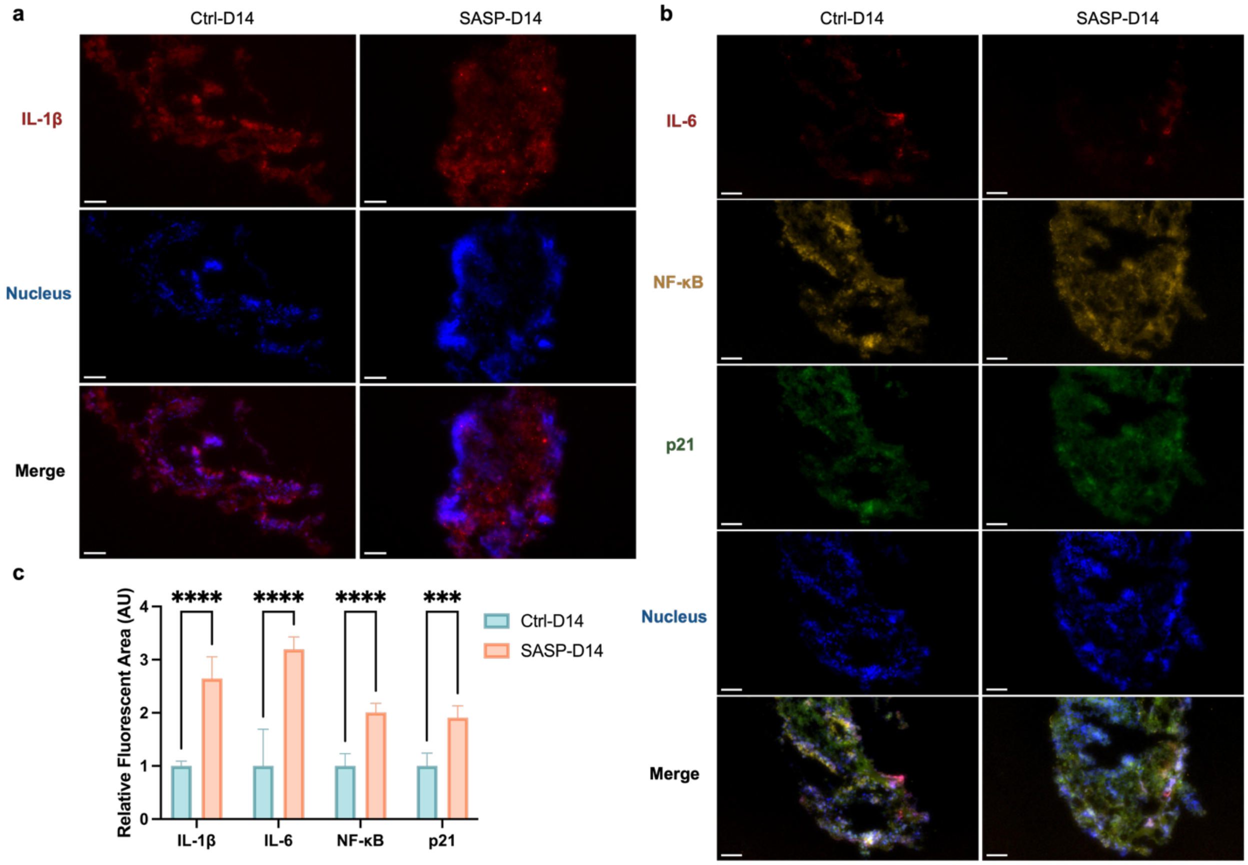


**Extended Data Fig. 3| Immunofluorescence analysis of cerebral organoids following SASP-CM treatment. a,** Representative immunofluorescence staining of cerebral organoids showing IL-1β (red) and nuclei (blue, DAPI). **b,** Representative multiplex immunofluorescence staining of cerebral organoids showing IL-6 (far-red), NF-κB (yellow), p21 (green), and nuclei (blue, DAPI). **c,** Quantification of fluorescence area for IL-1β, IL-6, NF-κB, and p21. Fluorescent regions were measured using Fiji (N=5). Values were normalized to the mean area of the Ctrl-D14 group for each marker. Data are shown as mean ± SD. Significance levels are denoted as ***: $p < 0.001$, ****: $p < 0.0001$. All Images in the figure were acquired using a 20× objective lens with a scale bar: 50μm
